# Light-independent and pulsing-sensitive magnetic compass responses in planarians

**DOI:** 10.64898/2026.09.27.754318

**Authors:** Leif Moritz, Evelyn Schmidt, Bénédicte Darbon, Aarya J. Pawar, Floriane Bureau, Pia Schürmann, Cigdem Ilayda Kalyon, Sybille Wolf-Kümmeth, Yannick Menger, Aude Chambodut, Hervé Cadiou, E. Pascal Malkemper

## Abstract

Many animals can sense the Earth’s magnetic field and use it for orientation. However, the mechanism underlying magnetoreception has not been identified, partially due to the lack of reproducible and genetically tractable model systems. Here, we establish a directional assay for planaria to test their responses to different magnetic fields. *Girardia dorotocephala* showed a preferred magnetic directionality in the horizontal plane in an Earth-strength magnetic field in the absence of visible light, which shifted if the azimuth was rotated and was absent when the magnetic field was cancelled. This directional preference differed between labs. Furthermore, the directional response was affected by a short but strong magnetic pulse, resulting in a random distribution. The effect of the pulse was temporary, with directional preferences recovering over several days. Taken together, these data are consistent with a magnetoreception mechanism in planarians that functions independently of light and relies on permanently magnetised material, such as single-domain magnetite. As planarians are genetically tractable and well-established, they are a promising invertebrate model organism that might offer insights into the mechanisms underlying magnetoreception in animals.

## INTRODUCTION

Magnetoreception, the ability to perceive the Earth’s magnetic field, has been demonstrated for representatives of various animal taxa (Lam and Malkemper, 2026), including arthropods (Ernst and Lohmann, 2016; Tenneriello, Wegmann and Fleischmann, 2026; Wang *et al*., 2026), molluscs (Sumner-Rooney *et al*., 2014), fish (Naisbett-Jones and Lohmann, 2022), birds (Wiltschko and Wiltschko, 2019) and mammals (Burda *et al*., 1990; Holland *et al*., 2008; Schneider *et al*., 2023). The geomagnetic field has a horizontal component, with field lines pointing toward magnetic North, and an inclination (i.e., the angle of the field lines toward Earth) and intensity that vary along the magnetic North-South axis. Intensity varies between ∼25 µT at the equator and ∼ 65 µT at the magnetic poles (Mouritsen, 2018). Thus, the geomagnetic field can serve as a reference system for local orientation and global navigation. Organisms can use different receptor mechanisms to detect these different properties of the geomagnetic field. However, the underlying mechanism remains unknown, and so far no magnetic field transducer has been identified in any organism other than magnetotactic bacteria (Nordmann, Hochstoeger and Keays, 2017). The most discussed putative magnetoreception mechanisms are a light-dependent radical pair mechanism, electromagnetic induction by movement in the magnetic field, and a magnetic particle-based mechanism (Shaw *et al*., 2015; Nordmann, Hochstoeger and Keays, 2017). Understanding of the underlying mechanism would be facilitated by a robust, tractable model system, ideally an invertebrate, for ethical and practical reasons. However, experiments demonstrating magnetoreceptive abilities in invertebrate standard models were often not reproducible and are still heavily debated, as in *Caenorhabditis elegans* (Vidal-Gadea et al., 2015; Malkemper et al., 2023) and *Drosophila melanogaster* (Bae *et al*., 2016; Bassetto *et al*., 2023).

Planarians are flatworms found in marine, freshwater, and terrestrial environments. Several freshwater planarian species are well-established model organisms in regeneration and developmental biology (Pagán, 2017), neuroscience (Benita, Nesher and Shomrat, 2024), and sensory systems research (Pearl, 1903; Inoue *et al*., 2015; Reho, Lelièvre and Cadiou, 2022; Sabry *et al*., 2022; Reho *et al*., 2024; Ascunce *et al*., 2026). First evidence for magnetoreception in freshwater planarians (Tricladida) was presented in a series of pioneering studies in the 1960s (Brown, 1962, 1965; Brown and Park, 1965b, 1965a; Brown and Chow, 1975) which showed that the mean direction in which the animals moved depended on the horizontal component of the magnetic field (azimuth), even comparing them to a “magnetic compass needle” (Brown, 1965). Despite these promising results, planarians received surprisingly little attention in magnetoreception research. Recent studies, however, have shown the effects of magnetic fields on regeneration (Jenrow, Smith and Liboff, 1996; Ermakov *et al*., 2022) and the modulation of ROS levels (Kinsey, Van Huizen and Beane, 2023; Rishabh *et al*., 2026) in planarians, but these effects are not necessarily specific and related to magnetoreception as a sensory modality.

Here we aimed to re-assess Brown’s initial experiments (Brown, 1962) in light of the advances in the field and test whether planarians orient with respect to Earth-strength magnetic fields. First and in order to set a reliable animal model of magnetoreception, we performed the experiments independently in two laboratories (INCI (HC) and MPINB (EPM)). We then aimed to gain initial insights into the underlying physical mechanism by testing in the absence of visible light and after magnetic pulsing, a diagnostic test for a magnetoreception mechanism based on magnetite (Kirschvink, Walker and Diebel, 2001).

## MATERIAL AND METHODS

### Study organism and husbandry

All tests were performed with asexually reproducing *Girardia dorotocephala* (Woodworth, 1897) (sometimes referred to as *Dugesia dorotocephala*), the same species that Brown (Brown, 1962) had demonstrated to react to weak magnetic fields (Fig. 1A). Animals were initially obtained from Carolina Biologicals (Burlington, North Carolina, US, GPS coordinates: 36°5′23″N 79°26′44″W, magnetic parameters in 2015: declination: −9° 02’ 53” inclination: 62° 46’ 41”, horizontal component: 22.4 µT, vertical component: 43,5 µT, total field: 48,9 µT) in 2015 and kept in the laboratory at INCI since. Specimens were transferred to the MPINB in Bonn in 2021. Animals were maintained in Volvic water at a 12/12 hour light-dark-circle (at MPINB 07:00 - 19:00 CET; at INCI 12:00 – 12:00 CET) within a wooden cabinet at MPINB at RT and in an electrically grounded incubator (Pol-Eko Apparatura, Poland) at 21°C at INCI. Animals were fed once a week with beef liver and starved for at least 7 days before experiments. Individuals were handled in dim red light (SOLRAOX 12V LED Strip red, 620 nm), which has been shown not to disturb the animals (Paskin *et al*., 2014), with a plastic Pasteur pipette with a cut-off and burned tip to avoid stressing and injuring the animals. Experiments started at the earliest 2 hours after the end of the light phase.

**Figure 1:**
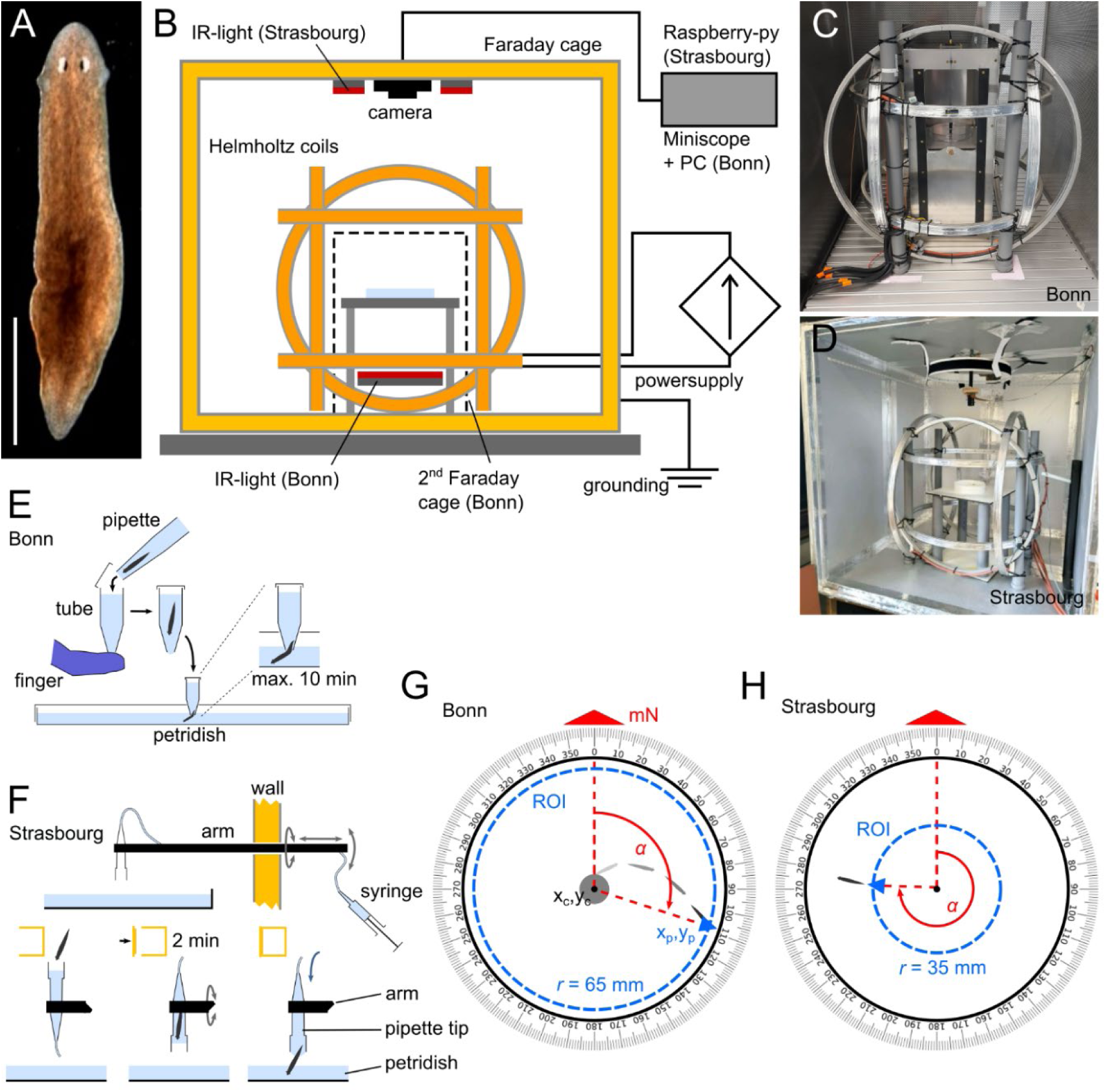
Planarians were tested in different magnetic fields and tracked to score their directional preference. **A** The freshwater planarian *Girardia dorotocephala* (Woodworth, 1897), which was reported by Brown (1962) to be capable of magnetoreception. Dorsal view. Scale = 1 mm. **B** Animals were tested in a circular arena (Petri dish) in the centre of a triple-wrapped three-axis Helmholtz coil system (blue), which allowed cancellation of the ambient magnetic field and creation of Earth-strength magnetic fields of different azimuths. The arena and coil were shielded by Faraday cages, and experiments were done under infrared illumination. C & D Similar Helmholtz coil systems were used in Bonn (C) and in Strasbourg (D). **E** Schematic representation of the passive release mechanism used in Bonn. **F** Schematic representation of the active release mechanism used in Strasbourg. **G** Schematic representation of the automated tracking in Bonn. The directional preference was scored 65 mm from the centre. **H** Schematic representation of the manual tracking in Strasbourg. Animals were tracked by hand, and the directional preference was scored 35 mm from the centre.

### Experiment 1: Directional assay - Bonn

Experiments at MPINB in Bonn (Germany) were carried out inside a triple-wrapped three-axis Helmholtz coil system (Claricent, Germany) placed inside a grounded Faraday cage (Figs 1B, C). For a detailed description of the setup, see Malkemper et al. (2023). The coil was powered by two low-residual-ripple Rohde & Schwarz HAMEG power supplies (HMP4030 & HMP4040). To expose the animals to different magnetic fields, a near-zero field (B <100 nT) was created with a set of compensation coils (first wrap), and on top of this near-zero field, magnetic fields were created with variations in the horizontal component. Azimuth relative to the back of the experimental chamber was set to 0° (mN-0: B ≈ 49 µT, D ≈ 0°, I ≈ 66°) or to 180° (mN-180: B ≈ 49 µT, D ≈ 180°, I ≈ 66°), thus rotating the local magnetic meridian using the additional two wraps of the coils. As is common in animal magnetoreception research, we here refer to the direction of the horizontal component of the magnetic field as magnetic North (mN). The field coils were set to antiparallel in the near-zero field condition (B < 100 nT), ensuring identical current flow and side effects such as heat or vibrations. We changed the magnetic field between runs to minimise daytime effects, and measured the field with an FVM-400 magnetometer (MEDA) at the start and end of each test day. The planarians were tested inside a second smaller aluminium Faraday cage placed in the Helmholtz coil on a non-magnetic stage that was levelled and centred in all three axes (Fig. 1B, C). The stage was illuminated from below by 12 V Synergy 21 infrared LED flex strips (S21-LED-F00107) (940 nm), and videos were recorded from above with a Miniscope MiniCAM V.3.3 camera (Open-ephys, OE-7451), with an 8 mm Arducam camera lens (M2508ZH02) with the infrared filter removed. Infrared light was used as planarians show no photophobic response to it (Paskin *et al*., 2014). For each run of the directional assay, a clean 145 mm plastic Petri dish was placed in the centre of the stage, filled with 80 ml of Volvic water, and covered with a lid with a centrally cut-out hole. A single individual was placed inside a black 1.5 ml Eppendorf tube with a cut tip (bottom). Each planaria was only tested once in one of the three conditions. After closing the tube, it was placed in the hole at the center of the Petri dish so that the tip was submerged in water, while the tested magnetic field was on. This way, the animal could exit the release device voluntarily at any time (Fig. 1E). The centre Faraday cage was closed, and the recording started. The recording lasted for 10 minutes after the animal left the release device or until it reached the margin of the Petri dish. If the individual did not leave the release device within 5 minutes, the run was stopped. Data was recorded in Bonsai 2.7.2 4 with a resolution of 2591×1944 pixels at 30 fps and saved as an mp4 file. From March 2023 to September 2024, the experiments were performed by three different experimenters. For logistical reasons (fire safety), the setup had to be moved within the same room between experiments.

Post-processing of videos and tracking of planarians was done in Python version 3.13.7. Initial videos recorded in Bonsai were resized with a custom Python script (ResizeVideoLoop_LM.py) to 800 x 600 pixels and saved as an AVI file to increase processing speed.

Tracking of individual movement was performed automatically using a custom Python script (TrackingVideoLoop_LM.py), which detected the centre of the arena and each individual in each frame via background subtraction, with a size- and intensity-based filter to avoid false detections of artifacts/noise during recording. For each frame, the x and y coordinates and the distance and angle relative to the centre were recorded. The angle at the point where the individual left a circular ROI with a radius of 65 mm (266 pixels) around the centre was scored as an individual’s directional preference (Figs 1G). We chose this radius based on pilot experiments (data not shown) and to maximise the radius (to allow the animal to orient while moving) while avoiding border effects near the Petri dish margin (r = 70 mm). After tracking, tracks were manually inspected by an experimenter blinded to the magnetic condition, and tracks were discarded if animals were not properly tracked or did not leave the ROI. The directional preference from all valid trials was extracted to a single Excel file in Python (Extract_rows_LM.py).

### Experiment 2: Directional assay - Strasbourg

To test for reproducibility of results, a slightly modified version of the experiment was carried out by independent experimenters at INCI in Strasbourg (France), where a possible magnetosensitive behaviour was first observed (unpublished data).

Experiments were carried out in Strasbourg, France between January 2025 and August 2026 (GPS: 48° 34’ 24.259” N 7° 45’ 7.601” E, magnetic parameters: declination: 3° 25’ 2”, inclination: 64° 32’ 5”, horizontal component 20.8 µT, vertical component: 43.9 µT, total field: 48.6 µT (September 2026). The laboratory is located at the South end of the first floor and is mostly surrounded by offices. This place was chosen for its low level of electromagnetic interference and the stability of the ambient magnetic field (although the horizontal component was approx. 4 µT lower than the predicted values).

The magnetic orientation set-up consisted of a cubic home-made light-tight Faraday cage connected to the ground (width: 100 cm, height: 100 cm, depth: 100 cm). In essence, panels of aluminium covered with insulated material (double-faced, thickness 1 mm) were assembled in order to get electrical conduction between all faces using aluminium tape (Fig. 1B, D). The test arena, a 14.5 cm glass Petri dish (with no markings, sitting on a 14.5 cm diameter white filter paper disc and surrounded by a 2 cm high circular white filter paper band), was placed in the centre of a 3D Helmholtz coils system (diameter 64 cm, Claricent, Germany) identical to the coil system used in Bonn. Each coil was double wrapped and connected to the ground. Double wrapping allowed the use of a sham condition with current flowing in anti-parallel directions. The coils were driven by computer-controlled (software: ETC, M631 « arbitrary waveform generator », Slovakia) power supplies (Kepco Inc., Korea). The inside of the Faraday cage was illuminated by a circular (31 cm Ø) infrared (850 nm) LED strip (BuyLEDstrip, Netherlands), allowing the infrared-sensitive camera (Arducam IMX477) to capture the area where worms were set. A 32 cm-tall stand was set in the centre to elevate the experimental zone 40 cm below the camera, which was attached to the ceiling of the chamber. The camera was controlled by a micro-computer (Raspberry Pi 4) outside the chamber. Videos were acquired at 10 frames per second. Before each experimental session, the magnetic field was measured with a 3-axis magnetometer (Metrolab, Switzerland) and the 3D coil system was realigned to magnetic north.

For the experiments, only large intact animals were selected. A single animal was introduced into the release device (Fig. 1F), a P1000 pipette tip mounted on a 1 m long PVC tube, and was left for 2 minutes in the closed Faraday cage. A gentle hydrostatic pressure was then applied in order to expel the worm at the centre of the arena (Fig. 1F). The worm was left in the experimental arena for two minutes and was subsequently taken out. Worms that failed to reach the virtual circular line of 35 mm radius around the centre within the two minutes were excluded from analysis. In between trials, the arena was thoroughly washed with ethanol rinsed 2–3 times (x2-3) with MilliQ water and 1 time with Volvic mineral water (x1) in order to get rid of possible chemical cues.

Animals were tested either in the natural ambient magnetic field in Strasbourg (B = 46– 48 µT), in a sham near-zero field (same setting as near-zero-field but current running antiparallel; 46–48 µT) or in a near-zero field (< 1 µT). In the latter two conditions, the same individuals were tested with the conditions in random order.

Videos were analysed using Tracker (Physlet, https://physlets.org/tracker/). The resulting tracks were then imported into GraphPad Prism (GraphPad software, Boston, MA, US), and a virtual circular line was drawn at a distance of 35 mm from the centre, and the angle where the track and the line met was reported in relation to the magnetic north (Fig. 1H) as the individual’s directional preference.

### Experiment 3: Electromagnetic pulsing

To test for the effect of electromagnetic pulsing, and the potential presence of magnetite-based magnetoreceptors, the directional preference of the planarians was tested before and after pulsing over consecutive days at INCI in Strasbourg (France). We constructed a custom pulsing device at MPINB, consisting of a double-wrapped Lee-Whiting four-coil system with a 90 mm diameter powered by a custom-built circuit (for design and a part list, see supplemental data). The pulse magnetizer circuit can generate a short, strong current pulse that drives the magnetic coils. Two capacitors, each with a capacitance of 390 µF, are charged to the set voltage. The voltage is converted from the 24 V input voltage to up to 120 V via a DPX800S CNC Booster module. The pulse can be triggered via a BNC connector, a screw terminal (galvanically isolated) or a pushbutton. Upon receiving an input signal, a silicon-controlled rectifier (MCR25N) triggers current flow from the pre-charged capacitors to the coils. A timing element ensures the capacitors can discharge completely and that the MCR25N switches off again. The pulser produced a ∼20 mT magnetic pulse with a current of 60V, when the coil pairs were set in parallel. The pulse intensity was measured with a F.W. Bell 5189 Gauss/Tesla Meter in the centre of the pulsing coils. The pulser was placed in the centre of a set of single-wrapped Helmholtz coils that created a 90 µT biasing field antiparallel to the pulsing direction, to ensure that potential magnetic particles within the animals are aligned opposite to the pulsing direction. We verified the pulse effect by pulsing a standard hiking compass, placed in the centre of the pulser set to parallel, within the biasing field. This led to a permanent flip of the polarity of the compass needle, pointing south instead of north in the ambient Earth-strength magnetic field. A subsequent pulse in the opposite direction reversed the effect.

For pulsing, we placed the animals in a petri dish without water for immobilisation. Animals were either exposed to a vertical biasing field (BF 90 µT) alone or a short vertical pulse (20 mT) antiparallel to the biasing field. We used a vertical pulse to orient all animals comparably relative to the pulse, because, as benthic animals, they mainly move and bend in the horizontal plane and barely in the vertical plane.

We tested the same animals in an Earth-strength magnetic field (47–48 µT) before the pulse, directly after pulsing, 1 day after pulsing, 3 days after pulsing, and 7 days after pulsing. The control group was tested before, right after and 1 day after the control treatment.

### Statistical analysis

After tracking and scoring each individual’s directional preference in a magnetic field, the group response in each magnetic field was tested using circular statistics. We performed statistical tests on the full Bonn dataset, including all subsets, to test the three conditions: mN-0, mN-180, and near-zero. The experimenter performing the tracking and statistical tests was blinded to which group corresponded to which condition. To test for a significant directional preference in each condition, a Rayleigh test was performed in Oriana version 4.02 (KCS). For the directional assays performed in Bonn, no correction for multiple testing was done, as independent groups of animals were tested under different conditions to test pre-defined hypotheses. This is common practice in the field (i.e. we could not find a single study testing directionality in a circular arena via Rayleigh test, which corrected for multiple testing). For the sham-near-zero field vs near-zero-field experiment and the pulsing experiments performed in Strasbourg, p-values were Holm-Bonferroni-adjusted (pcor) for multiple testing within sham-near-zero field vs near-zero-field animals as well as within the pulsed and control groups, as the same individuals were tested under different conditions. To test for significant differences in the distribution between the conditions, a pairwise Watson-U^2^ test was performed initially in Oriana. Oriana reports significance intervals (e.g. p < 0.05, < 0.01, < 0.001) and does not compute exact p-values. The same is true for the watson.two.test() implemented in the R package circular, and significance after correction for multiple testing could not be determined for all comparisons based on these approaches. Therefore, a permutation-based Watson-U^2^ test was performed in R (R Core Team, 2025) with the package circular (Agostinelli and Lund, 2025) and the function watson.two.test() (Ruxton, Malkemper and Landler, 2023). Based on the computed U^2^-scores of each pair, permutation-based p-values (100,000 simulations) were derived, followed by Holm-Bonferroni correction for multiple testing. To compute p-values from permutations, pairs were pooled, and 100,000 random samples of the original sample sizes were drawn without replacement from the pooled data. The U^2^-score was computed for each simulation, and the number of simulations with at least the original U^2^-score + 1 was divided by the number of simulations. For the pulsing experiments, a Watson-U^2^ test was performed only with comparison to the before-treatment group with 100,000 replicates and Holm-Bonferroni correction for multiple testing.

## RESULTS

### Planarians can sense an Earth-strength magnetic field in the absence of visible light (Experiments 1 & 2)

The tested planarians showed a directional preference in an earth-strength (B ≈ 49 µT) magnetic field under IR-light in the absence of visible light in two independent laboratories (Fig. 2). The animals tested in Bonn (Experiment 1) showed a significant directional group preference towards magnetic South in the mN-0-field (Rayleigh test: n = 131, µ = 187.0°, r = 0.2265, p = 0.0012, Fig. 2A, Tab. S1), and a significant directionality towards magnetic East in the rotated mN-180-field (Rayleigh test: n = 134, µ = 284.3°, r = 0.1906, p = 0.0055, Fig 2B, Tab. S1), while there was no directional preference in the near-zero-field (B < 100 nT; Rayleigh test: n = 127, µ = 102.5°, r = 0.0922, p = 0.3395, Fig. 2C, Tab. S1).

**Figure 2:**
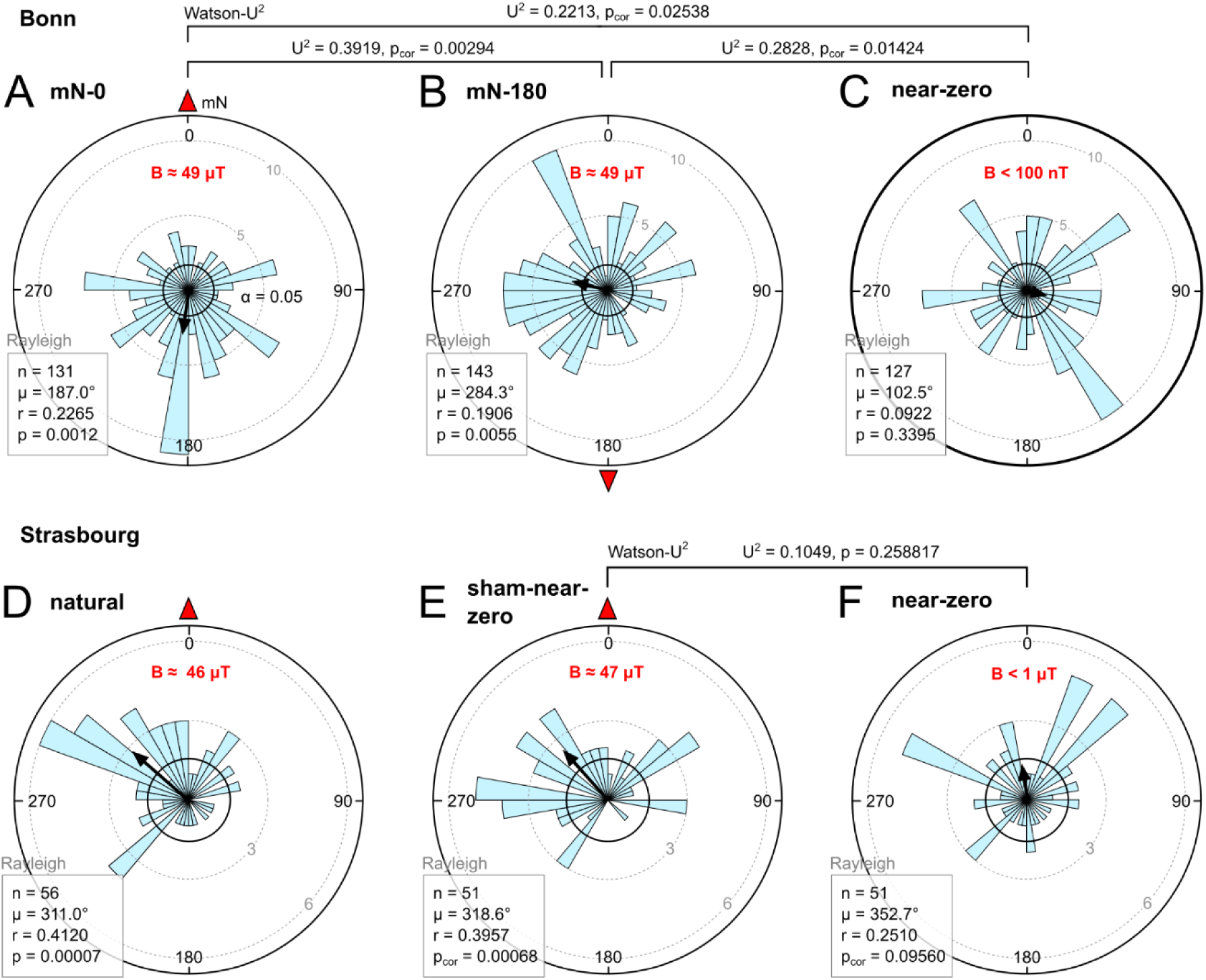
Planarians showed a directional preference in Earth-strength magnetic fields, but not in a near-zero magnetic field. Results of the directionality assay under different magnetic fields at MPINB (Bonn) and INCI (Strasbourg). **A – C** Experiments in Bonn, including all sets of experiments (n = 401). **A** mN-0, an Earth-strength field (49 µT) with an inclination of 66° and azimuth at 0°. **B** mN-180, an Earth-strength field (49 µT) with an inclination of 66° and azimuth at 180°. **C** near-zero, magnetic field reduced to <100 nT. **D – F** Experiments in Strasbourg. **D** Natural ambient field of Strasbourg. **E** Sham-near-zero field, with coils run with same currents as in F but antiparallel. **F** near-zero field. Magnetic north (mN) is indicated by a red arrowhead. Circular histograms show the number of individuals with a directional preference in 10° bins. The central arrow indicates the mean vector of the directional preference; if the group directionality differed significantly (Rayleigh p < 0.05) from a random/uniform distribution, the arrow surpasses the inner circle (95% significance interval). Sample size (n), mean vector (µ), and Rayleigh p-value (p) are given in each plot; For E and F, p-values were Holm-Bonferroni adjusted, as the same individuals were tested under different conditions. Pairwise Watson-U^2^ U^2^-scores (U^2^) and permutation (100,000) based Holm-Bonferroni-corrected p-values (p_cor_) are given above significance brackets (see also Tabs S1, S2). For results of individual sets, see Fig. S1 and Tab. S1.

A pairwise Watson-U^2^ test with permutation-based p-values and Holm-Bonferroni-correction showed significant differences between all conditions (mN-0 vs. mN-180: U^2^ = 0.3920, p_cor_ = 0.00294; mN-0 vs near-zero: U^2^ = 0.2213, p_cor_ = 0.02538; mN-180 vs near-zero: U^2^ = 0.2828, p_cor_ = 0.01424; Fig. 2, Tab. S2).

Independent experiments at INCI in Strasbourg (Experiment 2) in the natural ambient magnetic field showed a significant directional group-preference for magnetic North-West (Rayleigh test: n = 56, µ = 311.0°, r = 0.4120, p = 0.0000745; Fig. 2D, Tab. S3). While animals exposed to a sham-near-zero field, with current running through the coils but the ambient natural field maintained, showed a significant preference for magnetic North-West as well (Rayleigh test: n = 51, µ = 318.6°, r = 0.3957, p_cor_ = 0.00068; Fig. 2E, Tab. S3), animals exposed to a near-zero magnetic field showed a random distribution (Rayleigh test: n = 51, µ = 352.7°, r = 0.2510, p_cor_ = 0.09560; Fig. 2F, Tab. S3).

### Electromagnetic pulsing affects directionality (Experiment 3)

To test for a potential involvement of magnetic particle-based receptors, we performed pulsing experiments at INCI in Strasbourg. We found that the pulsing affected the orientation of the planarians, which showed a significant directional preference towards North-West before the treatment in the pulsed group (Rayleigh test: n = 20, µ = 313.0°, r = 0.7399, p_cor_ = 1.47×10^-5^, Fig. 3A, Tab. S4) and in the control group (Rayleigh test: n = 20, µ = 323.7°, r = 0.599, p_cor_ = 0.008, Fig. 3F, Tab. S4). Pulsing led to a random distribution (Rayleigh test: n = 20, µ = 350.5°, r = 0.3411, p_cor_ = 0.0967, Fig. 3B, Tab. S4), significantly different from the distribution before the pulse (Watson-U^2^: U^2^ = 0.3744, p_cor_ = 0.00268, Fig. 3, Tab. S4), an effect not observed in the control group (Rayleigh test: n = 20, µ = 339.4°, r = 0.5853, p_cor_ = 0.006, Fig. 3G, Tab. S4; Waton-U^2^: U^2^ = 0.0544, p_cor_ = 1, Fig. 3, Tab S4). Pulsed animals showed again a significant directionality already one day after the pulse (Rayleigh test: n = 20, µ = 23.3°, r = 0.4299, p_cor_ = 0.0456, Fig. 3C, Tab. S4), although the direction was significantly different from the original preference (Watson-U^2^: U^2^ = 0.3024, p_cor_ = 0.01206, Fig. 3, Tab. S4). Seven days after pulsing, the animals showed the original directional preference towards North-West (Rayleigh test: n = 20, µ = 304.6°, r = 0.5210, p_cor_ = 0.01, Fig. 3E, Tab. S4; Watson-U^2^: U^2^ = 0.1125, p_cor_ = 0.223688, Fig. 3, Tab S4).

**Figure 3:**
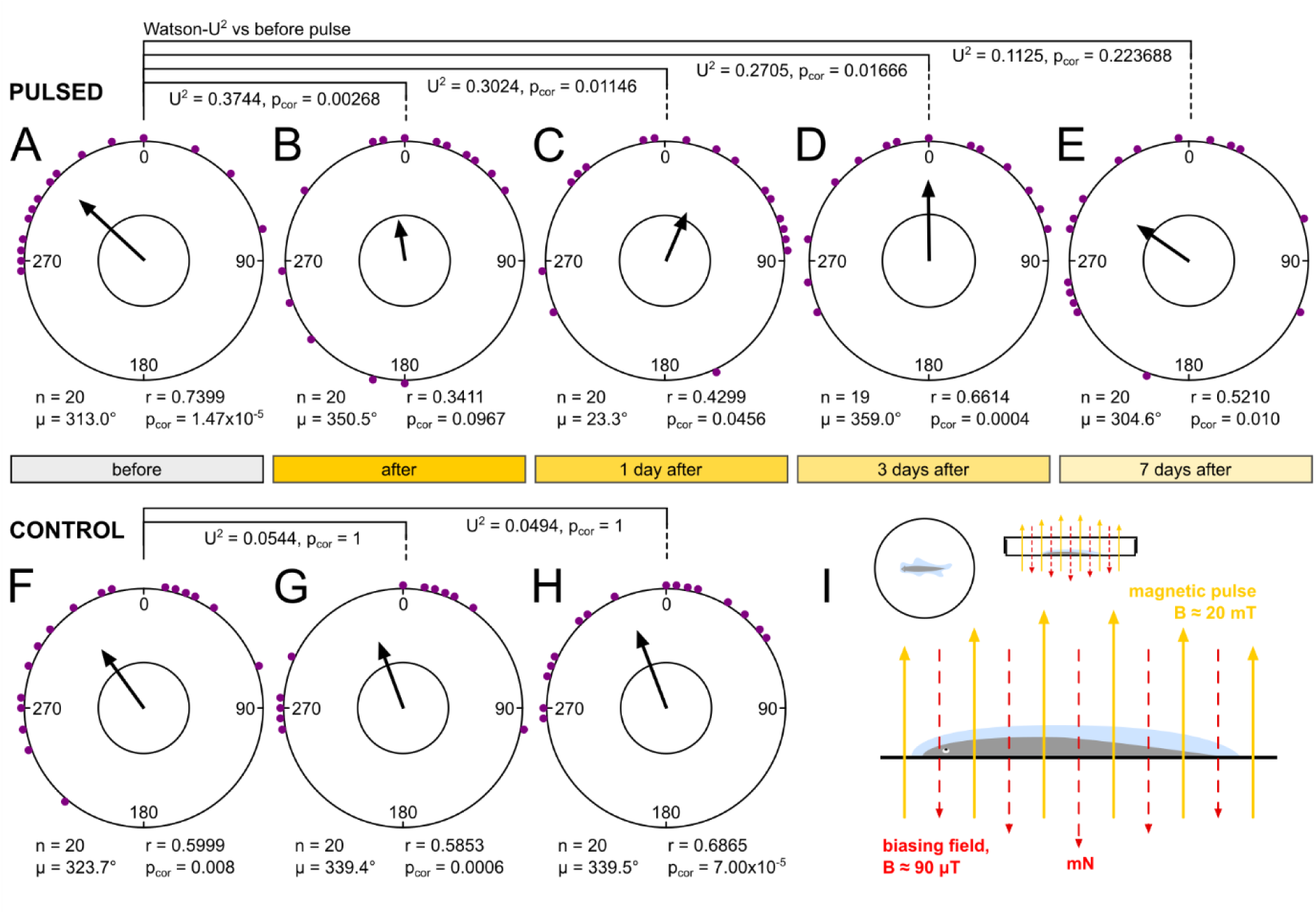
A short magnetic pulse temporarily disrupts magnetic orientation in *G. dorotocephala*. mN is in all trials at 0°; directionality was scored at a radius of 35 mm. **A – E** Directional preference of the pulsed group before (A), directly after (B), 1 day after (C), 3 days after (D) and 7 days after (E) exposure to a 20 mT magnetic pulse. **F – H** Directional preference of the control group before (F), directly after (G) and 1 day after (H) the control treatment. Circular plots show the directional preference of each individual. The central arrow indicates the mean vector of the directional preference. If the group directionality differed significantly (Rayleigh p < 0.05) from a random/uniform distribution, the arrow surpasses the inner circle. Sample size (n), mean vector (µ), and the Holm-Bonferroni-adjusted Rayleigh p-value (p_cor_) are shown below the plots (see also Tab. S4). Holm-Bonferroni adjustment was done within each group. A pairwise Watson-U2 test was performed to test the distribution in each trial against the distribution in the corresponding before-treatment trial. U^2^-scores (U2) and permutation (100,000) based Holm-Bonferroni-corrected p-values (p_cor_) are given under significance brackets (see also Tabs S4). **I** Treatment of the pulsed group. Animals were immobilised in a Petri dish by removing water around them, and a vertical 20 mT pulse was applied antiparallel to a 90 µT biasing field with magnetic North directed to the bottom.

## DISCUSSION

### Planarians can sense an Earth-strength magnetic field

The behavioural experiments in manipulated magnetic fields show that *G. dorotocephala* possess a magnetic compass sense, as the preference shifted with magnetic field rotations and was absent in a near-zero field, confirming Brown’s initial studies (Brown, 1962). Similar evidence has been presented for various other animals considered to have a magnetic sense (Wiltschko *et al*., 2008). Although the effect of the rotation of the Earth’s magnetic field on the group directionality was significant, surprisingly, the directionality changed by only 97.3° clockwise when magnetic North was rotated by 180°. This might be due to competing stimuli other than the magnetic field that affect their overall directionality, as planarians can sense a wide range of environmental stimuli (Pearl, 1903; Paskin *et al*., 2014; Inoue *et al*., 2015; Sabry *et al*., 2022; Ascunce *et al*., 2026). Surprisingly, magnetic directional preference differed between populations tested in Bonn and Strasbourg, suggesting that magnetic directional preference is not fixed but entrained or context-dependent.

### Behavioural data are congruent with a magnetite-based mechanism

Animals may perceive magnetic fields via different mechanisms (Johnsen and Lohmann, 2005; Nordmann, Hochstoeger and Keays, 2017). Behavioural data allow us to exclude certain mechanisms. Here we observe a magnetic response in the absence of visible light, as animals were tested under infrared light and handled under dim red light. This renders a radical pair mechanism for magnetoreception, as proposed for birds (Ritz, Adem and Schulten, 2000), unlikely because it is assumed to require visible, short-wavelength UV to green light (Wiltschko *et al*., 1993; Munro *et al*., 1997; Wiltschko and Wiltschko, 1999; Wiltschko, Nießner and Wiltschko, 2021; Xu *et al*., 2021).

Planarians exhibit electro- or galvanotaxis (Pearl, 1903; Viaud, 1954; Sabry *et al*., 2022), which in theory could enable them to sense the geomagnetic field via electromagnetic induction. However, we consider an induction-based magnetoreception, as proposed for rays and sharks and recently also pigeons (Kalmijn, 1971, 1982; Nimpf *et al*., 2019; Nordmann *et al*., 2026), to be unlikely in planarians because of their small size, their slow movement of 1-5 mm/s (Reho, Lelièvre and Cadiou, 2022), their low behavioral sensitivity to electric field of ca. 0.33 V/cm (Sabry *et al*., 2022), compared to 5 nV/cm in sharks (Kalmijn, 1971, 1982; Nimpf *et al*., 2019), and because the studied species dwells in a weakly electric conducting media i.e. freshwater.

As a radical-pair-and an induction-based mechanism are unlikely, magnetoreception in planarians might be based on magnetic particles, i.e. magnetite particles associated with membranes or mechanosensitive channels (Nordmann, Hochstoeger and Keays, 2017). A diagnostic test for the involvement of such particles has been to expose animals to a short but strong pulse in the mT range. Magnetite is expected to be permanently remagnetized by such a magnetic pulse, altering its behaviour in the magnetic field and the response of potential magnetite-based receptors (Kirschvink, Walker and Diebel, 2001; Davila *et al*., 2005; Johnsen and Lohmann, 2005; Shaw *et al*., 2015). An effect of electromagnetic pulsing on the orientation of animals has been demonstrated for birds (Wiltschko *et al*., 2006; Holland and Helm, 2013), turtles (Mackiewicz *et al*., 2025), fish (Naisbett-Jones *et al*., 2020), mammals (Marhold, Wiltschko and Burda, 1997; Holland *et al*., 2008), and spiny lobsters (Ernst and Lohmann, 2016), leading to a change or absence of the directional preferences (Johnsen and Lohmann, 2005). In our experiments, pulsing led to a loss of a significant magnetic directional preference in planarians. These results support the presence of a magnetite-based receptor, and future studies should focus on identifying candidate magnetoreceptors by screening for single-domain magnetite particles (Moritz *et al*., 2026), which would be sufficient to open mechanosensitive channels (Winklhofer and Kirschvink, 2010). Interestingly, a significant directionality returned within one day after pulsing and the animals showed the original directional preference towards North-West a week after pulsing. This is consistent with experiments on migratory birds (*Zosterops lateralis*), in which a return to their original heading was observed after 10 days (Wiltschko *et al*., 2006).

### Limitations and outlook

Our data suggest that planarian magnetic responses are weak and can be easily masked by other environmental factors and stimuli more relevant to the animals. The mean directionality varied between sessions (Fig. S1, Tab. S1), and relatively large sample sizes were required for a significant group response in the Malkemper lab (MPINB, Bonn). Furthermore, the rotation of the directional preference did not fully reach the magnetic field rotation (97.3° vs 180°), pointing to competing stimuli that might enhance the magnetic directional preference in mN-0 and counteract it in mN-180. Furthermore, in subsequent trials at different locations within the laboratory (MPINB, Bonn), we observed a significant directionality independent of the applied magnetic fields. Magnetoreception might be disturbed by other external (fluctuating) influences (e.g., magnetic radiofrequencies, electric fields) or overpowered by stimuli higher in the hierarchy (e.g., light, vibration, temperature). In some birds, magnetoreception is mainly relied upon in the absence of visual cues (Zein *et al*., 2026), and experiments testing the hierarchy of sensory modalities in planarians did not include magnetic fields (Inoue *et al*., 2015). Experimenters may not be aware of these stimuli outside their own perception, or they may be hard to control in a laboratory setting, such as electromagnetic fields, radiofrequencies, or low-frequency vibrations.

Because we studied the group response, and animals were tested only once, individuals’ navigational abilities and directional preferences could not be tested. In birds, it has been shown that not all individuals show the same motivation to orient in the magnetic field, and that pre-selecting motivated animals can improve the results (Patmanidis *et al*., 2026). Further work on planarians should address this question by taking advantage of this peculiar animal model and by selecting clones based on their orientation and magnetic selectivity.

The biological relevance of a magnetic sense in planarians in their natural environment remains unknown and can only be speculated. The magnetic directional preference might be entrained to avoid aversive stimuli, like light or certain surface textures. In previous studies, it has been shown that planarians can learn to avoid or find certain locations (Best and Rubinstein, 1962; Jordan *et al*., 2023; Qian *et al*., 2023) and to associate certain contexts with negative stimuli via classical conditioning (Thompson and McConnell, 1955; James *et al*., 2026). The entrainment of a preferred magnetic directionality is supported by the different orientations observed in the populations from Bonn and Strasbourg. Magnetoreception in planarians might serve in finding their way back to suitable shelter after foraging as a homing mechanism or, as it has been suggested for some animals (Bae *et al*., 2016), might serve as a gravitational sense, which could also be the case in planarians since these animals do not sense gravity through specialised cells but rather through non-specific cellular mechanisms (Adell *et al*., 2014).

## Conclusion

We here present data from two laboratories that support a magnetic sense in planarians, as initially demonstrated by Brown in the 60’s (Brown, 1962). This mechanism is likely to be mediated by magnetite, suggesting them as a promising, genetically tractable invertebrate model system for the magnetic sense. Planarians are easy to handle and maintain in the lab (Merryman, Alvarado and Jenkin, 2018). They reproduce via fission and have remarkable regenerative abilities, which enable amputation experiments to further determine the position of receptors (Paskin *et al*., 2014; Inoue *et al*., 2015; Sabry *et al*., 2022). Furthermore, genes can be knocked down via RNA interference (Sánchez Alvarado and Newmark, 1999 refined by Reho *et al*., 2025). We therefore encourage replication experiments on directional responses in additional labs and hope that future investigations use the planarian toolkit to narrow down the magnetoreceptive cells and shed light on the transduction mechanism.

## Supporting information

Supplemental data (magnetic pulser)

## AKNOWLEDGEMENTS

We thank the NMR group at MPINB for discussions and support during the research project. AJP obtained and analysed data in the course of the M.Sc. OEP Biology at the University of Bonn. ES obtained and analysed data in the course of the B.Sc. Applied Biology at the Hochschule Bonn-Rhein-Sieg. We are further indebted to Bernd Scheiding at the MPINB electronics workshop for constructing the magnetic pulser. We thank Pierrick Poisbeau and his team for initial discussions about the project. We are grateful to Etienne Challet/Valérie Simonneaux and their team for invaluable advice.

## USE OF AI

ChatGPT (OpenAI) GPT-5.6 Luna was used to assist with the generation, debugging, and refinement of data-analysis scripts. All code was reviewed and validated by the authors, who take full responsibility for the analyses and reported results. No AI was used for the writing of the manuscript.

## AUTHORS CONTRIBUTION

Conceptualisation: HC, LM, EPM. Formal Analysis: LM, AJP, ES, PS. Funding Acquisition: HC, EPM. Investigation: LM, AJP, ES, PS, SWK, BD, HC, FB, CIK, YM. Methodology: All authors. Resources: HC, EPM. Validation: LM, EPM, HC. Visualisation: LM. Writing – Original Draft Preparation: LM, EPM. Writing – Review & Editing: LM, EPM, HC. Technical support: FB, YM.

## DATA AVAILABILITY

The data generated in the study as well as scripts used for analysis are available upon reasonable request from the corresponding authors.

## SUPPLEMENTARY MATERIAL

### Supplementary figures

**Figure S1:**
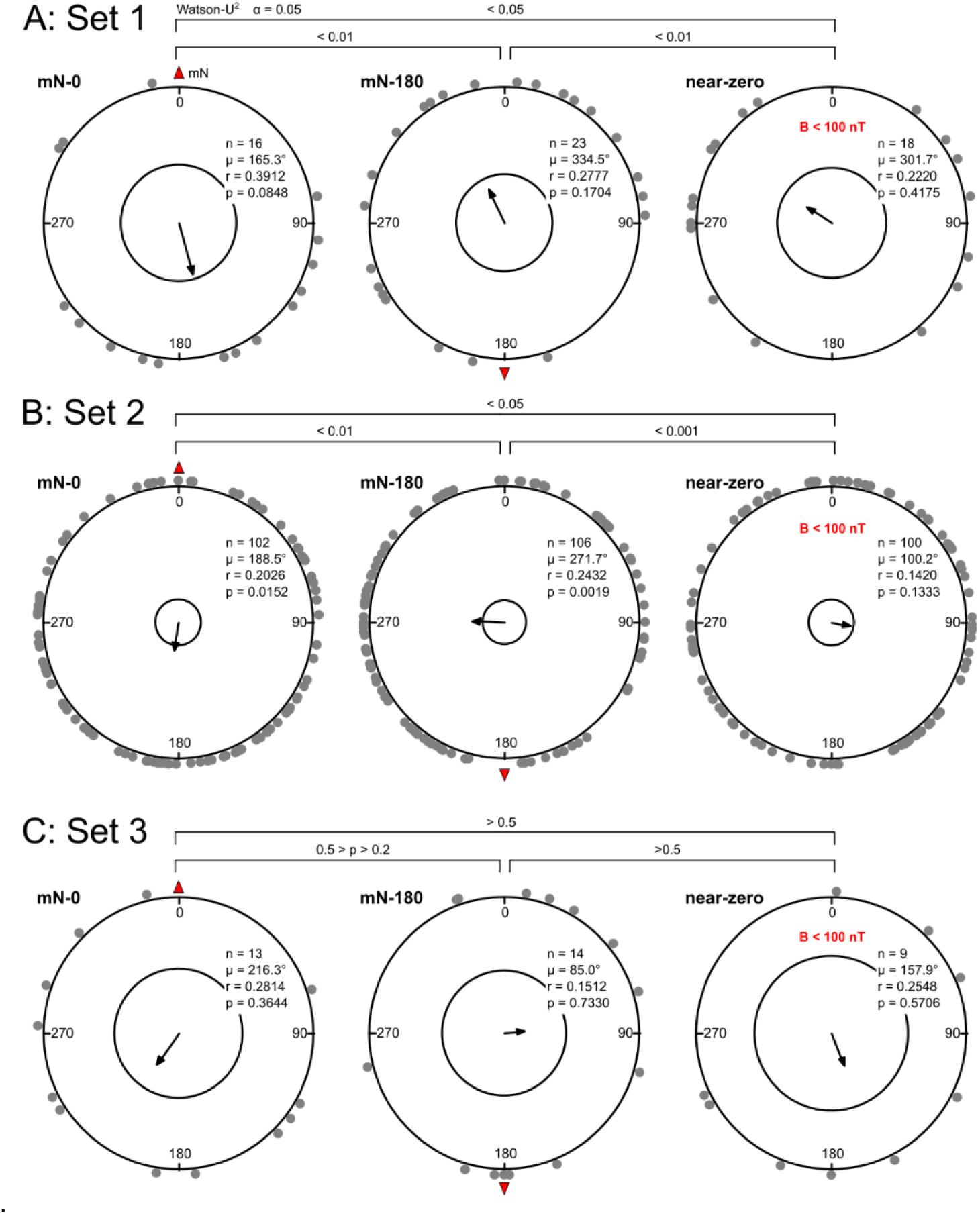
Planarians’ directional preference shows high variability. Results of the directionality assay under different magnetic fields in the three individual sets of experiments (n = 401) performed by different experimenters. **A** Set 1, 27.03.2023–06.04.2023 by AJP. **B** Set 2, 05.12.2023–23.01.2024 by ES. **C** Set 3, 25. & 26.09.2024 by SWK. Each dot around the outer circle represents the directional preference of a single individual tested once. The central arrow indicates the mean vector of the directional preference; if the group directionality differed significantly (p < 0.05) from a random/uniform distribution, the arrow surpasses the inner circle (α = 0.05). Sample size (n), mean vector (µ) and Rayleigh p-value (p) are given within the plots. Pairwise Watson-U^2^ p-values are given above significance brackets.

### Supplementary tables

**Table S1:** Statistical analysis, including Rayleigh test for each condition and experimental set tested at MPINB in Bonn (Germany), Experiment 1. Tests were performed in Oriana, and results were rounded to 4 decimals. The p-values (p) of the total dataset were adjusted for multiple testing using Holm-Bonferroni correction (p_cor_). “All” shows the results for the combined data. cond. = condition, n = number of individuals, µ = mean vector (°), r = length pf mean vector, conc. = concentration, var = circular variance, SD = circular standard deviation, z = Rayleigh z, p = Rayleigh p, p_cor_ = p-values after Holm-Bonferroni-correction for multiple testing.

| Cond. | mN-0 |  |  |  | mN-180 |  |  |  | near-zero |  |  |  |
| --- | --- | --- | --- | --- | --- | --- | --- | --- | --- | --- | --- | --- |
| Set | Set1 | Set2 | Set3 | All | Set1 | Set2 | Set3 | All | Set1 | Set2 | Set3 | All |
| n | 16 | 102 | 13 | 131 | 23 | 106 | 14 | 143 | 18 | 100 | 9 | 127 |
| $\mu$ | 165.32 | 188.47 | 216.30 | 187.01 | 334.45 | 271.73 | 84.98 | 284.28 | 301.69 | 100.15 | 157.88 | 102.46 |
| r | 0.3912 | 0.2026 | 0.2814 | 0.2265 | 0.2777 | 0.2432 | 0.1512 | 0.1906 | 0.2220 | 0.1420 | 0.2548 | 0.0922 |
| conc. | 0.8499 | 0.4138 | 0.3242 | 0.4650 | 0.5782 | 0.5016 | 0.0000 | 0.3884 | 0.4555 | 0.2868 | 0.1055 | 0.1852 |
| var. | 0.6088 | 0.7974 | 0.7186 | 0.7735 | 0.7223 | 0.7568 | 0.8488 | 0.8094 | 0.7780 | 0.8580 | 0.7452 | 0.9078 |
| SD | 78.50 | 102.38 | 91.24 | 98.75 | 91.71 | 96.34 | 111.36 | 104.32 | 99.40 | 113.21 | 94.74 | 125.09 |
| z | 2.4488 | 4.1872 | 1.0293 | 6.7182 | 1.7739 | 6.2719 | 0.3202 | 5.1970 | 0.8874 | 2.0152 | 0.5844 | 1.0802 |
| p | 0.0848 | 0.0152 | 0.3644 | 0.0012 | 0.1704 | 0.0019 | 0.7330 | 0.0055 | 0.4175 | 0.1333 | 0.5706 | 0.3395 |
| $p_{\text{cor}}$ | | | | 0.0036 | | | | 0.0110 | | | | 0.3395 |

**Table S2:** Results of the pairwise Watson-U^2^ test to compare the distribution between conditions tested at MPINB in Bonn (Germany), Experiment 1. The Watson-U^2^ test was performed in Oriana, and additionally U^2^-scores were calculated in the circular package (Agostinelli and Lund, 2025) in R (R Core Team, 2025). In R, p-values (p) were derived from permutations (100,000), and a Holm-Bonferroni correction for multiple testing was performed (p_cor_) in R. Note that Oriana gives significance intervals but not discrete p-values.

| Group1 | Group2 | Oriana |  | Permutation based |  |  |
| --- | --- | --- | --- | --- | --- | --- |
|  |  | U<sup>2</sup> | p | U<sup>2</sup> | p | p<sub>cor</sub> |
| mN-0 | mN-180 | 0.391777 | < 0.001 | 0.391901 | 0.00098 | 0.00294 |
| mN-0 | zero | 0.220853 | < 0.05 | 0.221341 | 0.02538 | 0.02538 |
| mN-180 | zero | 0.282817 | < 0.01 | 0.282823 | 0.00712 | 0.01424 |

**Table S3:**
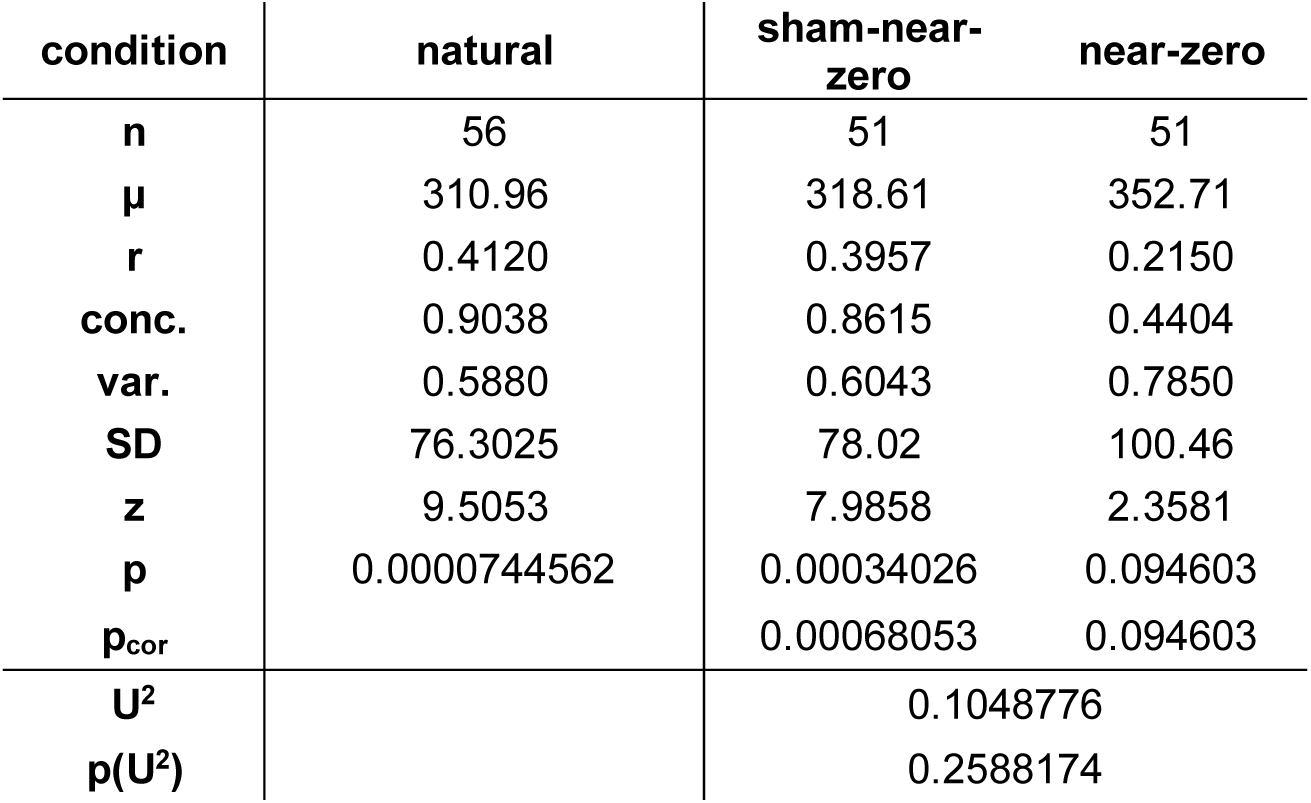
Results of the Rayleigh test and the pairwise Watson-U^2^ to compare the distribution between conditions tested at INCI in Strasbourg (France), Experiment 2. We performed the Rayleigh test in Oriana, and rounded results to 4 decimals. The p-values (p) of the sham-near-zero and near-zero treatment were adjusted for multiple testing using Holm-Bonferroni correction (p_cor_), as the same individuals were tested. For the Watson-U^2^-test between sham-near-zero and near-zero, U^2^-scores were calculated in the circular package (Agostinelli and Lund, 2025) in R (R Core Team, 2025). In R, p-values (p) were derived from permutations (100,000). n = number of individuals, µ = mean vector (°), r = length pf mean vector, conc. = concentration, var = circular variance, SD = circular standard deviation, z = Rayleigh z, p = Rayleigh p, p_cor_ = p-values after Holm-Bonferroni-correction for multiple testing, U^2^ = Watson-U^2^-score, p(U^2^) = permutation based p-value of Watson-U2-test.

**Table S4:** Statistical analysis, including Rayleigh test for the pulsed and control groups tested at INCI in Strasbourg (France), Experiment 3. The groups were tested before treatment (BP), directly after treatment (AP) and 1 day (1DAP), 3 days (3DAP) and 7 days after treatment (7DAP). Statistical tests were performed in Oriana. The p-values of the pulsed (5 tests) and control (3 tests) datasets were corrected separately for multiple testing using Holm-Bonferroni correction. For the Watson-U^2^-test against the before-pulse condition (BF) within each experiment, U^2^-scores were calculated in the circular package (Agostinelli and Lund, 2025) in R (R Core Team, 2025). p-values (p) were derived from permutations (100,000), and a Holm-Bonferroni correction for multiple testing was performed (p_cor_) in R. cond. = condition, n = number of individuals, µ = mean vector (°), r = length of mean vector, conc. = concentration, var = circular variance, SD = circular standard deviation (°), z = Rayleigh z, p = Rayleigh p, p_cor_ = p-values after Holm-Bonferroni-correction for multiple testing, U^2^ = Watson-U^2^-score, p(U^2^) = permutation based p-value of Watson-U2-test, p_cor_(U^2^) = permutation based p-value of Watson-U2-test with Holm-Bonferroni correction for multiple testing.

| experiment<br>condition | Pulsed |  |  |  |  | control |  |  |
| --- | --- | --- | --- | --- | --- | --- | --- | --- |
|  | BF | AP | 1DAP | 4DAP | 7DAP | BF | AF | 1DAF |
| n | 20 | 20 | 20 | 19 | 20 | 20 | 20 | 20 |
| $\mu$ | 312.96 | 350.54 | 23.28 | 359.02 | 304.64 | 323.70 | 339.42 | 339.53 |
| r | 0.7399 | 0.3411 | 0.4299 | 0.6614 | 0.5210 | 0.5999 | 0.5853 | 0.6865 |
| conc. | 2.2817 | 0.7257 | 0.9514 | 1.7892 | 1.2155 | 1.5084 | 1.4506 | 1.9256 |
| var. | 0.2601 | 0.6589 | 0.5701 | 0.3386 | 0.4790 | 0.4001 | 0.4147 | 0.3135 |
| SD | 44.47 | 84.03 | 74.45 | 52.10 | 65.42 | 57.93 | 59.30 | 49.70 |
| z | 10.9493 | 2.3270 | 3.6954 | 8.3109 | 5.4297 | 7.1967 | 6.8525 | 9.4246 |
| p | 2.9364E-06 | 0.09663231 | 0.0227959 | 9.56251E-05 | 0.0033419 | 0.000415 | 0.0006306 | 2.33471E-05 |
| p<sub>cor</sub> | 1.4682E-05 | 0.09663231 | 0.0455918 | 0.0003825 | 0.0100256 | 0.00083 | 0.0006306 | 7.00413E-05 |
| U<sup>2</sup> |  | 0.3744375 | 0.302438 | 0.27049033 | 0.1125 |  | 0.0544375 | 0.0494375 |
| p(U<sup>2</sup>) |  | 0.00067 | 0.00382 | 0.00832992 | 0.223688 |  | 0.700713 | 0.75547245 |
| p<sub>cor</sub>(U<sup>2</sup>) |  | 0.00268 | 0.01146 | 0.01665983 | 0.223688 |  | 1 | 1 |

## REFERENCES

Adell, T. et al. (2014) “Planarians Sense Simulated Microgravity and Hypergravity,” BioMed Research International, 2014(1), p. 679672. Available at: 10.1155/2014/679672.

Agostinelli, C. and Lund, U. (2025) “R package ‘circular’: Circular Statistics (version 0.5-2).” Available at: https://CRAN.R-project.org/package=circular.

Ascunce, K. et al. (2026) “The Sensory Basis of Planarian Behavior,” Journal of Neuroscience, 46(13). Available at: 10.1523/JNEUROSCI.1158-25.2025.

Bae, J.-E. et al. (2016) “Positive geotactic behaviors induced by geomagnetic field in Drosophila,” Molecular Brain, 9(1), p. 55. Available at: 10.1186/s13041-016-0235-1.

Bassetto, M. et al. (2023) “No evidence for magnetic field effects on the behaviour of Drosophila,” Nature, 620(7974), pp. 595–599. Available at: 10.1038/s41586-023-06397-7.

Benita, O., Nesher, N. and Shomrat, T. (2024) “Neurophysiological measurements of planarian brain activity: a unique model for neuroscience research,” Biology Open, 13(8), p. bio060480. Available at: 10.1242/bio.060480.

Best, J.B. and Rubinstein, I. (1962) “Maze learning and associated behavior in planaria,” Journal of Comparative and Physiological Psychology, 55(4), pp. 560–566. Available at: 10.1037/h0044394.

Brown, F.A. (1962) “Responses of the planarian, dugesia, and the protozoan, paramecium, to very weak horizontal magnetic fields,” The Biological Bulletin, 123(2), pp. 264–281. Available at: 10.2307/1539273.

Brown, F.A. (1965) “A unified theory for biological rythms,” Circadian Clocks - Proceedings of the Feldafing Summer School 7-18 September 1964. Amsterdan: North-Holland Publishing Company.

Brown, F.A. and Chow, C.S. (1975) “Differentiation between Clockwise and Counterclockwise Magnetic Rotation by the Planarian, Dugesia dorotacephala,” Physiological Zoology, 48(2), pp. 168–176. Available at: 10.1086/physzool.48.2.30155650.

Brown, F.A. and Park, Y.H. (1965a) “Duration of an after-effect in planarians following a reversed horizontal magnetic vector,” The Biological Bulletin, 128(3), pp. 347–355. Available at: 10.2307/1539898.

Brown, F.A. and Park, Y.H. (1965b) “Phase-shifting a lunar rhythm in planarians by altering the horizontal magnetic vector,” The Biological Bulletin, 129(1), pp. 79–86. Available at: 10.2307/1539768.

Burda, H. et al. (1990) “Magnetic compass orientation in the subterranean rodentCryptomys hottentotus (Bathyergidae),” Experientia, 46(5), pp. 528–530. Available at: 10.1007/BF01954256.

Davila, A.F. et al. (2005) “Magnetic Pulse Affects a Putative Magnetoreceptor Mechanism,” Biophysical Journal, 89(1), pp. 56–63. Available at: 10.1529/biophysj.104.049346.

Ermakov, A. et al. (2022) “Effect of weak alternating magnetic fields on planarian regeneration,” Biochemical and Biophysical Research Communications, 592, pp. 7–12. Available at: 10.1016/j.bbrc.2021.12.096.

Ernst, D.A. and Lohmann, K.J. (2016) “Effect of magnetic pulses on Caribbean spiny lobsters: implications for magnetoreception,” Journal of Experimental Biology, 219(12), pp. 1827–1832. Available at: 10.1242/jeb.136036.

Holland, R.A. et al. (2008) “Bats Use Magnetite to Detect the Earth’s Magnetic Field,” PLOS ONE, 3(2), p. e1676. Available at: 10.1371/journal.pone.0001676.

Holland, R.A. and Helm, B. (2013) “A strong magnetic pulse affects the precision of departure direction of naturally migrating adult but not juvenile birds,” Journal of the Royal Society Interface, 10(81), p. 20121047. Available at: 10.1098/rsif.2012.1047.

Inoue, T. et al. (2015) “Planarian shows decision-making behavior in response to multiple stimuli by integrative brain function,” Zoological Letters, 1(1), p. 7. Available at: 10.1186/s40851-014-0010-z.

James, F.L. et al. (2026) “Adapting rodent cued threat conditioning to planarians: Memory acquisition, consolidation, and reconsolidation.” Available at: https://research-repository.rmit.edu.au/articles/journal_contribution/Adapting_rodent_cued_threat_conditioning_to_planarians_Memory_acquisition_consolidation_and_reconsolidation/31524397/1 (Accessed: July 9, 2026).

Jenrow, K. a., Smith, C. h. and Liboff, A. r. (1996) “Weak extremely-low-frequency magnetic field-induced regeneration anomalies in the planarian Dugesia tigrina,” Bioelectromagnetics, 17(6), pp. 467–474. Available at: 10.1002/(SICI)1521-186X(1996)17:6%3C467::AID-BEM6%3E3.0.CO;2-1.

Johnsen, S. and Lohmann, K.J. (2005) “The physics and neurobiology of magnetoreception,” Nature Reviews Neuroscience, 6(9), pp. 703–712. Available at: 10.1038/nrn1745.

Jordan, L. et al. (2023) “Conditioned place avoidance in the planaria *Schmidtea mediterranea*: A pre-clinical invertebrate model of anxiety-related disorders,” Behavioural Processes, 210, p. 104894. Available at: 10.1016/j.beproc.2023.104894.

Kalmijn, Ad.J. (1982) “Electric and Magnetic Field Detection in Elasmobranch Fishes,” Science, 218(4575), pp. 916–918. Available at: 10.1126/science.7134985.

Kalmijn, A.J. (1971) “The Electric Sense of Sharks and Rays,” Journal of Experimental Biology, 55(2), pp. 371–383. Available at: 10.1242/jeb.55.2.371.

Kinsey, L.J., Van Huizen, A.V. and Beane, W.S. (2023) “Weak magnetic fields modulate superoxide to control planarian regeneration,” Frontiers in Physics, 10. Available at: 10.3389/fphy.2022.1086809.

Kirschvink, J.L., Walker, M.M. and Diebel, C.E. (2001) “Magnetite-based magnetoreception,” Current Opinion in Neurobiology, 11(4), pp. 462–467. Available at: 10.1016/S0959-4388(00)00235-X.

Lam, B.Y. and Malkemper, E.P. (2026) “Magnetosensation-the unsolved mystery,” Physiological Reviews [Preprint].

Mackiewicz, A.G. et al. (2025) “Disruption of the sea turtle magnetic map sense by a magnetic pulse,” Journal of Experimental Biology, 228(22), p. jeb251243. Available at: 10.1242/jeb.251243.

Marhold, S., Wiltschko, W. and Burda, H. (1997) “A Magnetic Polarity Compass for Direction Finding in a Subterranean Mammal,” Naturwissenschaften, 84(9), pp. 421–423. Available at: 10.1007/s001140050422.

Merryman, M.S., Alvarado, A.S. and Jenkin, J.C. (2018) “Culturing Planarians in the Laboratory,” in J.C. Rink (ed.) Planarian Regeneration: Methods and Protocols. New York, NY: Springer, pp. 241–258. Available at: 10.1007/978-1-4939-7802-1_5.

Moritz, L. et al. (2026) “Multimodal imaging reveals no evidence for magnetite-based magnetoreceptors in the mole-rat eye.” bioRxiv, p. 2026.04.02.715577. Available at: 10.64898/2026.04.02.715577.

Mouritsen, H. (2018) “Long-distance navigation and magnetoreception in migratory animals,” Nature, 558(7708), pp. 50–59. Available at: 10.1038/s41586-018-0176-1.

Munro, U. et al. (1997) “Effect of Wavelength of Light and Pulse Magnetisation on Different Magnetoreception Systems in a Migratory Bird,” Australian Journal of Zoology, 45(2), pp. 189–198. Available at: 10.1071/ZO96066.

Naisbett-Jones, L.C. et al. (2020) “Magnetoreception in fishes: the effect of magnetic pulses on orientation of juvenile Pacific salmon,” Journal of Experimental Biology, 223(10), p. jeb222091.

Naisbett-Jones, L.C. and Lohmann, K.J. (2022) “Magnetoreception and magnetic navigation in fishes: a half century of discovery,” Journal of Comparative Physiology A, 208(1), pp. 19–40. Available at: 10.1007/s00359-021-01527-w.

Nimpf, S. et al. (2019) “A putative mechanism for magnetoreception by electromagnetic induction in the pigeon inner ear,” Current Biology, 29(23), pp. 4052–4059.

Nordmann, G.C. et al. (2026) “A global screen for magnetically induced neuronal activity in the pigeon brain,” Science, 391(6790), pp. 1155–1160. Available at: 10.1126/science.aea6425.

Nordmann, G.C., Hochstoeger, T. and Keays, D.A. (2017) “Magnetoreception—A sense without a receptor,” PLOS Biology, 15(10), p. e2003234. Available at: 10.1371/journal.pbio.2003234.

Pagán, O. (2017) “Planaria: an animal model that integrates development, regeneration and pharmacology,” International Journal of Developmental Biology, 61(8–9), pp. 519–529. Available at: 10.1387/ijdb.160328op.

Paskin, T.R. et al. (2014) “Planarian Phototactic Assay Reveals Differential Behavioral Responses Based on Wavelength,” PLOS ONE, 9(12), p. e114708. Available at: 10.1371/journal.pone.0114708.

Patmanidis, I. et al. (2026) “Directionality range in Emlen funnels,” Journal of the Royal Society, Interface, 23(234), p. rsif.2025.0225. Available at: 10.1098/rsif.2025.0225.

Pearl, R. (1903) “The Movements and Reactions of Fresh-water Planarians: a Study in Animal Behaviour.1,” Journal of Cell Science, s2-46(184), pp. 509–714. Available at: 10.1242/jcs.s2-46.184.509.

Qian, R. et al. (2023) “Spatial localization ability of planarians identified through a light maze paradigm,” PLOS ONE, 18(7), p. e0288118. Available at: 10.1371/journal.pone.0288118.

R Core Team (2025) “R: A Language and Environment for Statistical Computing.” Available at: https://www.R-project.org/.

Reho, G. et al. (2024) “Behavioral and pharmacological characterization of planarian nociception,” Frontiers in Molecular Neuroscience, 17. Available at: 10.3389/fnmol.2024.1368009.

Reho, G. et al. (2025) “Planarian RNAi knockdown: feeding once might just be enough,” Frontiers in Neuroscience, 19. Available at: 10.3389/fnins.2025.1546196.

Reho, G., Lelièvre, V. and Cadiou, H. (2022) “Planarian nociception: Lessons from a scrunching flatworm,” Frontiers in Molecular Neuroscience, 15. Available at: 10.3389/fnmol.2022.935918.

Rishabh, R. et al. (2026) “Verification of quantum biological predictions for weak magnetic field effects on superoxide in planarians,” PNAS Nexus, 5(5), p. pgag152. Available at: 10.1093/pnasnexus/pgag152.

Ritz, T., Adem, S. and Schulten, K. (2000) “A Model for Photoreceptor-Based Magnetoreception in Birds,” Biophysical Journal, 78(2), pp. 707–718. Available at: 10.1016/S0006-3495(00)76629-X.

Ruxton, G.D., Malkemper, E.P. and Landler, L. (2023) “Evaluating the power of a recent method for comparing two circular distributions: an alternative to the Watson U2 test,” Scientific Reports, 13(1), p. 10007. Available at: 10.1038/s41598-023-36960-1.

Sabry, Z. et al. (2022) “Head removal enhances planarian electrotaxis,” Journal of Experimental Biology, 225(17), p. jeb243972. Available at: 10.1242/jeb.243972.

Sánchez Alvarado, A. and Newmark, P.A. (1999) “Double-stranded RNA specifically disrupts gene expression during planarian regeneration,” Proceedings of the National Academy of Sciences, 96(9), pp. 5049–5054. Available at: 10.1073/pnas.96.9.5049.

Schneider, W.T. et al. (2023) “Migratory bats are sensitive to magnetic inclination changes during the compass calibration period,” Biology Letters, 19(11), p. 20230181. Available at: 10.1098/rsbl.2023.0181.

Shaw, J. et al. (2015) “Magnetic particle-mediated magnetoreception,” Journal of The Royal Society Interface, 12(110), p. 20150499. Available at: 10.1098/rsif.2015.0499.

Sumner-Rooney, L.H. et al. (2014) “Do chitons have a compass? Evidence for magnetic sensitivity in Polyplacophora,” Journal of Natural History, 48(45–48), pp. 3033–3045. Available at: 10.1080/00222933.2014.959574.

Tenneriello, C., Wegmann, J.W. and Fleischmann, P.N. (2026) “Magnetoreception in Cataglyphis hellenica ants,” Journal of Comparative Physiology A, 212(3), pp. 517–526. Available at: 10.1007/s00359-026-01794-5.

Thompson, R. and McConnell, J. (1955) “Classical conditioning in the planarian, Dugesia dorotocephala,” Journal of Comparative and Physiological Psychology, 48, pp. 65–68. Available at: 10.1037/h0041147.

Viaud, G. (1954) “Conception nouvelle du galvanotropisme animal Expérience sur les Planaires,” Experientia, 10(6), pp. 233–242. Available at: 10.1007/BF02157383.

Wang, W. et al. (2026) “The role of neuronal protein NOS in regulation of magnetic orientation in the subterranean termite Odontotermes formosanus,” Insect Molecular Biology, n/a(n/a). Available at: 10.1111/imb.70060.

Wiltschko, R. et al. (2008) “Light-dependent magnetoreception: orientation behaviour of migratory birds under dim red light,” Journal of Experimental Biology, 211(20), pp. 3344–3350. Available at: 10.1242/jeb.020313.

Wiltschko, R., Nießner, C. and Wiltschko, W. (2021) “The Magnetic Compass of Birds: The Role of Cryptochrome,” Frontiers in Physiology, 12, p. 667000. Available at: 10.3389/fphys.2021.667000.

Wiltschko, R. and Wiltschko, W. (2019) “Magnetoreception in birds,” Journal of The Royal Society Interface, 16(158), p. 20190295. Available at: 10.1098/rsif.2019.0295.

Wiltschko, W. et al. (1993) “Red light disrupts magnetic orientation of migratory birds,” Nature, 364(6437), pp. 525–527. Available at: 10.1038/364525a0.

Wiltschko, W. et al. (2006) “Bird navigation: what type of information does the magnetite-based receptor provide?,” Proceedings of the Royal Society B: Biological Sciences, 273(1603), pp. 2815–2820. Available at: 10.1098/rspb.2006.3651.

Wiltschko, W. and Wiltschko, R. (1999) “The effect of yellow and blue light on magnetic compass orientation in European robins, Erithacus rubecula,” Journal of Comparative Physiology A, 184(3), pp. 295–299. Available at: 10.1007/s003590050327.

Winklhofer, M. and Kirschvink, J.L. (2010) “A quantitative assessment of torque-transducer models for magnetoreception,” Journal of The Royal Society Interface, 7(suppl_2), pp. S273–S289. Available at: 10.1098/rsif.2009.0435.focus.

Xu, J. et al. (2021) “Magnetic sensitivity of cryptochrome 4 from a migratory songbird,” Nature, 594(7864), pp. 535–540. Available at: 10.1038/s41586-021-03618-9.

Zein, B. et al. (2026) “Geomagnetic night vision: A data-driven approach to study geomagnetic navigation in a migratory bird,” Ecological Modelling, 516, p. 111560. Available at: 10.1016/j.ecolmodel.2026.111560.

