## Supplementary figures and images for "Light-independent and pulsing-sensitive magnetic compass responses in planarians"

### Pulser.pdf

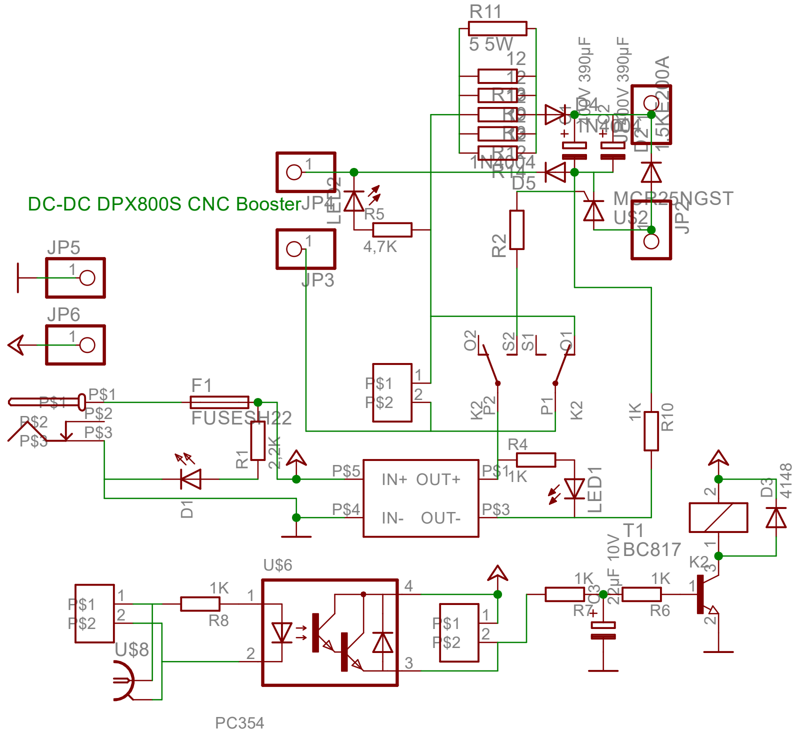

### Pulser_brd.pdf

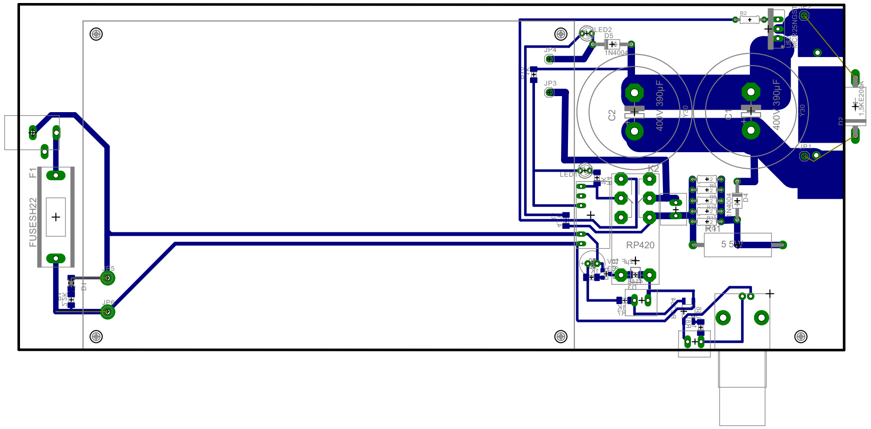

10.09.2026 09:37:59 f=0.50 Y:\E-Werkstatt\2024\E-Werk\Schaltungen\Pascal\Induki4\IndukiGrundPlatteklein.brd
